# The faecal metabolome and its determinants in inflammatory bowel disease

**DOI:** 10.1101/2022.06.15.495746

**Authors:** Arnau Vich Vila, Shixian Hu, Sergio Andreu-Sánchez, Valerie Collij, B. H. Jansen, Hannah E. Augustijn, Laura Bolte, Renate A.A.A. Ruigrok, Galeb Abu-Ali, Cosmas Giallourakis, Jessica Schneider, John Parkinson, Amal Al Garawi, Alexandra Zhernakova, Ranko Gacesa, Jingyuan Fu, Rinse K. Weersma

## Abstract

**Objective:** Inflammatory bowel disease (IBD) is a multifactorial immune-mediated inflammatory disease of the intestine, comprising Crohn’s disease and ulcerative colitis. By characterising metabolites in faeces, combined with faecal metagenomics, host genetics and clinical characteristics, we aimed to unravel metabolic alterations in IBD.

**Design:** We measured 1,684 different faecal metabolites and 8 short-chain and branched-chain fatty acids in stool samples of 424 IBD patients and 255 non-IBD controls. Regression analyses were used to compare concentrations of metabolites between cases and controls and determine the relationship between metabolites and each participant’s lifestyle, clinical characteristics and gut microbiota composition. Moreover, genome-wide association analysis was conducted on faecal metabolite levels.

**Results:** We identified over 300 molecules that were differentially abundant in the faeces of patients with IBD. The ratio between a sphingolipid and L-urobilin could discriminate between IBD and non-IBD samples (AUC = 0.85). We found changes in the bile acid pool in patients with dysbiotic microbial communities and a strong association between faecal metabolome and gut microbiota. For example, the abundance of *Ruminococcus gnavus* was positively associated with tryptamine levels. In addition, we found 158 associations between metabolites and dietary patterns, and polymorphisms near *NAT2* strongly associated with coffee metabolism.

**Conclusion:** In this large-scale analysis, we identified alterations in the metabolome of patients with IBD that are independent of commonly overlooked confounders such as diet and surgical history. Considering the influence of the microbiome on faecal metabolites, our results pave the way for future interventions targeting intestinal inflammation.

## INTRODUCTION

Characterisation of the host–microbiota symbiosis is crucial in the context of intestinal disorders such as inflammatory bowel disease (IBD) in which the gut environment is severely perturbed, yet the disease-causing mechanisms are still largely unknown. IBD is a chronic inflammatory disorder of the gastrointestinal tract that consists of two main subtypes: ulcerative colitis (UC) and Crohn’s disease (CD)^1,2^. In IBD, periods of active disease are characterised by loss of strictly anaerobic bacteria, blooming of facultative anaerobes and alterations in the chemical environment in the gut^3^. For example, reductions of gut barrier–protecting short-chain fatty acids (SCFA) and alterations in bile acids, sphingolipids and tryptophan-derived metabolites have been consistently reported in faeces of patients with IBD^4,5^. However, a large number of molecules in the human body remain uncharacterised, and thus their implications for human health remain unknown. Considering that a subset small molecules, including microbiome-derived metabolites, have been shown to regulate the immune response, it is crucial to characterise these metabolites and understand which factors determine their concentrations in the gut.

Recent technological advances in mass spectrometry techniques have enabled high-throughput characterisation and quantification of a wide range of known and chemically unannotated molecules^6^. In this context, the characterisation of faecal metabolites holds great potential for discovering non-invasive biomarkers and therapeutic targets. To date, however, studies performing untargeted metabolomics on the faeces of patients with IBD have been scarce, limited in sample size and lacking in in-depth information on host genetics, lifestyle, diet and clinical characteristics^4,7^.

In this study, we aimed to determine alterations in the gut metabolism of patients with IBD to pinpoint factors influencing faecal metabolite levels. For this, we used untargeted faecal metabolomics to characterise the prevalence and levels of 1,684 different faecal metabolites in a cohort of 424 patients with IBD and 255 controls. Our findings highlight the potential of faecal metabolites as biomarkers for IBD and show that, despite the influence of lifestyle, genetics and disease, faecal microbes are a strong predictor of the levels and composition of metabolites in the gut.

## METHODS

### Cohort and metadata description

Samples were obtained from two established cohorts: LifeLines^8^, a population biobank from the north of the Netherlands, and 1000IBD^9^, a cohort of patients with IBD from the University Medical Centre of Groningen (UMCG). In this study, we included 255 non-IBD controls, 238 patients with CD and 174 patients with UC. Sample collection, storage and cohort characteristics have been described before^10,11^ (**Suppl. Table 1**).

### Metabolite quantification

Metabolomics measurements performed by Metabolon Inc. (North Carolina, USA) (see Suppl. Methods) detected 1,684 different faecal metabolites (**Suppl. Table 2**). In addition to the untargeted metabolomics, we also measured the concentrations of eight short-chain and branched-chain fatty acids using liquid chromatography with tandem mass spectrometry (LC-MS/MS) methods **(Suppl. Table 3)**.

### Metabolic data processing

Metabolomic data was handled as a compositional dataset and transformed using centred log-ratios. Metabolites were split into three categories based on their prevalence in our cohort. The first group consisted of metabolites present in more than 70% of the samples in both the cases and controls (x = 854). Missing values were imputed using k-Nearest Neighbour Imputation using the “kNN” function from the “VIM” R package^12^. We set the number of nearest neighbours to 10 (k = 10) for the imputation and Euclidean distance as a metric. The second group of metabolites (prevalence < 70% and > 20%, x = 514) showed large sparsity across samples, so their levels were transformed into binary traits (metabolite presence/absence). Rare metabolites (prevalence < 20%, x = 316) were excluded from analysis given their limited power to detect meaningful associations.

### Identification of metabolites associated with IBD

To identify metabolites associated with IBD, we performed linear regression analysis using the *lm* function in R. The abundance of each metabolite was compared between disease groups (IBD/CD/UC) and controls. Technical factors (storage time, input grams of faeces and sample batch), host characteristics (age, sex, BMI and bowel movements per day), intestinal integrity (any resection: yes/no) and 24 dietary patterns that were significantly different between cases and controls were included as covariates in the regression models **(Suppl. Table 4)**. Less prevalent metabolites (prevalence < 70% and > 20% of samples) were evaluated using logistic regressions. Missing values were transformed to 0 and non-zero values to 1. Logistic regressions were performed using the implementation in the glm function in R and the covariates described above. All p-values were adjusted for multiple testing using Benjamini-Hochberg corrections as implemented in the *p.adjust* function. We considered a false discovery rate (FDR) < 0.05 as the threshold for statistical significance.

### Association between metabolites and phenotypes

An association analysis between phenotypes and metabolites was performed within each cohort (controls, CD and UC). We included information about lifestyle, including use of 31 different types of medication, dietary patterns represented by 144 food frequency–related scores and the levels of 3 faecal biomarkers (faecal calprotectin, chromogranin A and human beta-defensin) (see Suppl. Methods). Each phenotype– metabolite combination was tested using linear regression, including age, sex, BMI, bowel movements per day and technical factors as covariates.

### Co-occurrence patterns between bacteria and metabolites

*Mmvec* v.1.0.6^13^ was used to estimate the co-occurrence probabilities between highly prevalent metabolites and bacteria. The *QIIME*^14^ implementation of *mmvec* was used with default settings. Co-occurrence patterns were represented in a biplot using the ordination coordinates obtained from *mmvec* analysis. Next, we assessed the associations between individual microbiome features (taxa, gene clusters and metabolic pathways) and metabolites using regression models.

### Differential abundance analyses of faecal microbiome features

Linear regression analysis was used to identify microbiome features (taxa, pathways and metabolic gene clusters) that differed between controls and IBD. Age, sex, BMI, average bowel movements per day, history of intestinal resections (yes/no) and sequencing read depth were included as covariates in the regression models. Details of the processing of metagenomics data can be found in the Supplementary Methods.

### Association between dysbiosis and faecal metabolites

Phenotypic differences between patients with dysbiotic and eubiotic microbiota were established using chi-squared or Wilcoxon-rank test for categorical and continuous variables, respectively. Differences in the abundance of faecal metabolites between the two groups of patients were tested using linear regression. Age, sex, BMI, intestinal resection (yes/no), ileocecal valve *in-situ* (yes/no), average bowel movements per day and differences in 12 dietary patterns (**Suppl. Table 5**) were added as covariates in the regression models. Associations were considered statistically significant at FDR < 0.05.

### Meta-analysis of phenotype–metabolite and microbiome–metabolite associations

The results of the metabolite–phenotype and metabolite–microbiome analyses were combined in a meta-analysis using random-effects models implemented in R package *meta* (v.4.8). Results were considered statistically significant when the meta-FDR < 0.05.

### Genome-wide association analysis on faecal metabolites

Exome sequencing and genomic array data were available for both cohorts (see Suppl. Methods). Associations between host genetics and faecal metabolite levels were identified using regression models. Linear regression was used for metabolites present in > 70% of the samples and logistic regression for those present in between 70% to 20% of the samples. Analyses were performed per cohort, and results were combined in a meta-analysis, as previously described^15^. In addition to accounting for the confounders described above (see *Identification of metabolites associated with IBD*), we included population genetic structure as a covariate in the analysis. To determine the statistical significance of our findings we adopted two thresholds: a genome-wide significance (p-value<5e^−08^), and a more conservative cut-off, a study-wide significance (p-value <2.97e^−11^). The study-wide significance threshold was determined by dividing the genome-wide threshold by total number of metabolites (5.0e^−08^/1684).

### Prediction of IBD based on metabolomics profiles

We used CoDaCoRe^16^ (v 0.0.1) to identify ratios of metabolites and bacterial abundances that could predict IBD and its sub-phenotypes. Here, we first split the data into a training and a test set for each prediction, using 75% of the samples in the training process. Next, we estimated the added predictive value of using ratios of metabolites compared to a model including only host age, sex, BMI and faecal calprotectin levels (calprotectin levels >200 μg/g, yes/no). Patients with a history of intestinal surgeries (n = 136) were excluded, and only highly prevalent metabolites (>70% of the samples) were considered in this analysis.

### Metabolite level prediction

To predict the levels of metabolites in faeces, we performed regression models with L1 regularisation (lasso) using the glmnet R package^17^. We defined eight different models representing the data categories available in both cohorts (IBD and non-IBD samples, see Suppl. Methods). For each metabolite, we performed a 5-fold cross-validation (CV) procedure to select the best set of predictors based on the mean of squared errors. A 10-fold CV step was used in each of the CV-training sets to tune the lasso penalty parameter (lambda) in the lasso regression. Using the estimates of the model minimising the mean of squared errors, we computed the R^2^ coefficient in the whole dataset.

## RESULTS

### Patients with IBD have a distinct faecal metabolite profile

Metabolites were assessed in the faecal samples from 238 patients with CD, 174 patients with UC and 255 non-IBD controls. On average, 1,011 metabolites were detected per sample, ranging from 784 to 1,241 molecules. Patients with CD had a significantly larger number of metabolites in faeces than controls and patients with UC (Wilcoxon-test p-value< 2×10^−16^).

PCA based on the abundances of 854 highly prevalent metabolites revealed that IBD samples are dispersed across a cluster that partially overlaps with controls (Fig 1A). The first component of the PCA captured 18% of the variation and was driven by the levels of carnitine and bile and fatty acids, while the second component, representing 8% of cohort variation, was driven by the abundance of dipeptides and several unclassified metabolites (**Suppl. Table 6**, Fig 1B-C).

**Figure 1.**
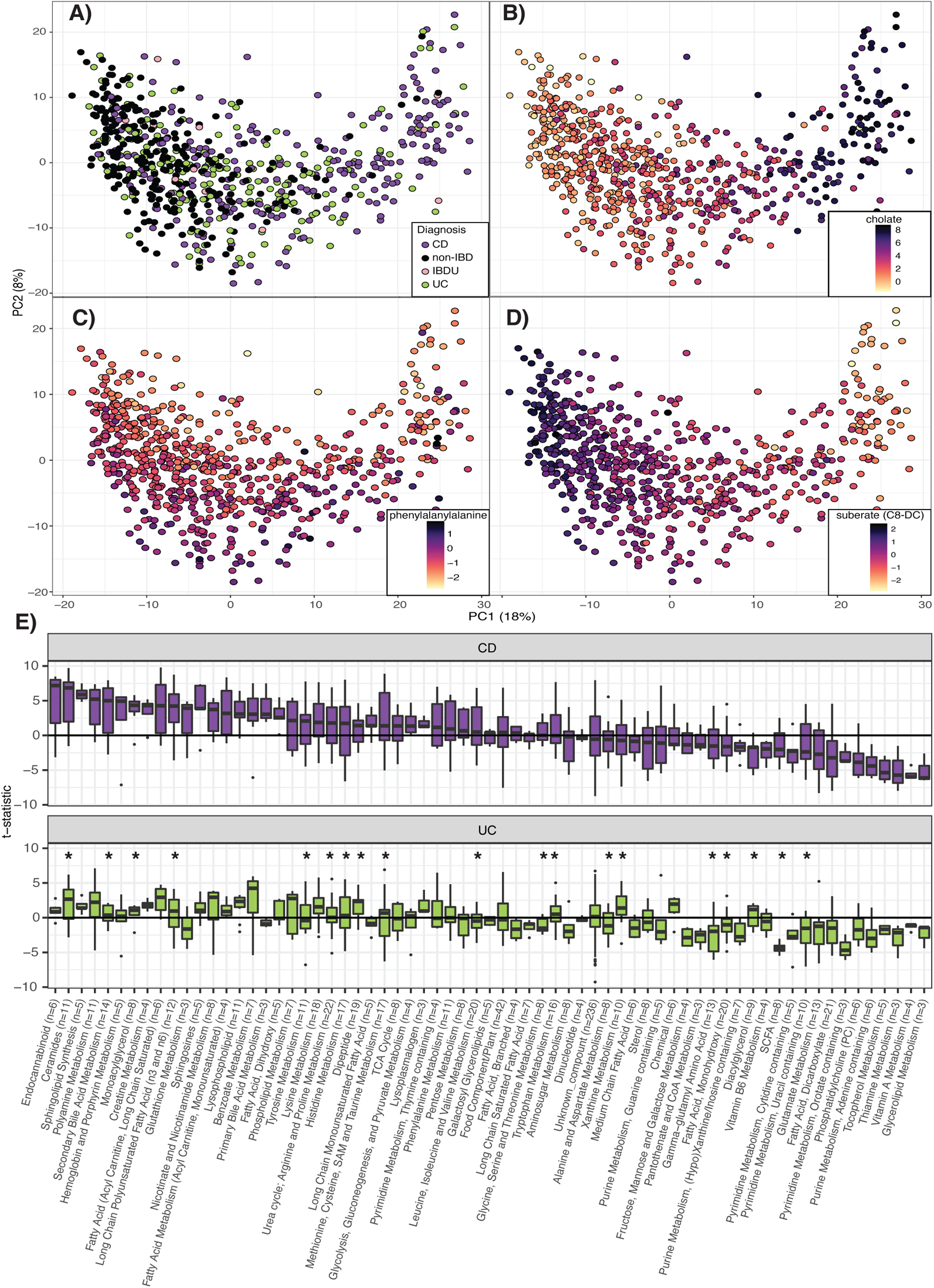
Faecal metabolite alterations in patients with Crohn’s disease and ulcerative colitis. **A-D.** Principal coordinate analyses depicting the clustering of 255 non-IBD (black), 238 CD (purple), 174 UC (green) and 12 IBDU (pink) samples according to their metabolomic composition. The first principal component is mainly driven by the levels of cholic acid and suberate (panel B and D) and the second component by the concentrations of phenylalanylalanine (panel C). Light– dark colour gradient represents low–high metabolite values. Metabolite concentrations are expressed as centred log-ratio (clr) of the AUC raw values. **E.** Metabolite differences between cases and controls grouped into metabolomic pathways. For clarity, only categories with three or more metabolites are shown (number of metabolites per categories are indicated on the x-axis). Y-axis represents the t-statistic value from the linear regression model (see Methods). Asterisk indicate significant differences between CD and UC (FDR < 0.05, Suppl. Table 7).

Differential abundance analysis revealed 324 associations when comparing patients with CD to non-IBD controls and 309 associations when comparing patients with UC to non-IBD controls (FDR<0.05) (**Suppl. Table 7,** **Suppl. Figure 1A**). Moreover, when looking into lower prevalence metabolites (present in < 70% of samples), we found that products of the metabolism of bile acids, ceramides and steroids were more prevalent in faeces of patients with IBD than in controls (182 and 119 molecules associated to CD and UC, respectively) (**Suppl. Table 7**).

A prominent signal in both disease groups was the depletion of vitamins and fatty acid– related molecules compared to controls (**Fig 1E**). In contrast, several derivatives of amino acid metabolism increased in the faeces of patients with IBD, which suggests an increase in amino acid utilisation in the gut of patients with IBD. For example, patients with IBD presented higher levels of the phenolic compound p-cresol sulphate, and multiple tryptophan derivatives from the indole/Ahr and kynurenine/IDO pathways were altered in IBD samples. The level of indole-propionic acid was decreased in UC (FDR_UC_ = 0.03), while tryptamine and kynurenine were increased in both CD and UC (FDR < 0.05). Patients with IBD also showed higher levels of arachidonic acid (20:4n6) and a lower ratio of omega-6/omega-3 fatty acids (**Suppl. Table 8, Suppl. Figure 1E**).

We also found that 106 metabolites were differentially abundant between CD and UC. For example, patients with UC presented higher levels of diaminopimelate (DAPA), an alpha-amino acid present in the cell membrane of gram-negative bacteria. Interestingly, DAPA-containing peptidoglycans can trigger the immune response mediated by *NOD1*^18^ (**Suppl. Figure 1C, Suppl. Table 9**).

### Patients with UC show the lowest concentrations of SCFAs in faeces

The SCFAs are a well-studied group of metabolites in the context of IBD. The concentrations of these molecules are essential for immune modulation, and their synthesis is dependent on colonic bacterial fermentation of polysaccharides^19^. Acetate, propionate and butyrate, the three most abundant SCFAs in the gut, were found in lower concentrations in patients with UC when compared with controls (FDR_UC_ < 0.05). Strikingly, after correcting for potential confounding effects, no significant differences in these metabolites were observed between CD and controls. In contrast, levels of hexanoic (or caproic acid) and valeric acids were significantly lower in both groups of patients. In addition, three branched-chain fatty acids (2-methylbutyrate, isobutyrate and isovaleric acid) were also found in lower concentrations in patients with UC (**Suppl. Figure 2, Suppl. Table 7**).

### Gut health and lifestyle are reflected in the levels of faecal metabolites

Besides the link between IBD and faecal metabolite composition, we assessed the association between faecal metabolites and 229 host characteristics, including dietary habits, medication use and clinical features such as inflammation location and history of intestinal resections. Due to the significant differences observed in metabolite composition between cases and controls, we carried out association analysis per condition (i.e. CD only, UC only and controls only) and then combined the results in a meta-analysis (**see Methods**).

Chromogranin A, beta-defensin 2 and calprotectin levels in faeces showed a large number of associations with faecal metabolites. In non-IBD controls, the level of chromogranin A was correlated with the levels of 164 metabolites, including positive associations with N-formylmethionine, cholesterol and secondary bile acids. Participants with calprotectin levels > 200 μg/g showed lower levels of cytidine in faeces. In addition, calprotectin was related to increased sphingosines and ceramides in UC but not CD (**Fig 2A**).

**Figure 2.**
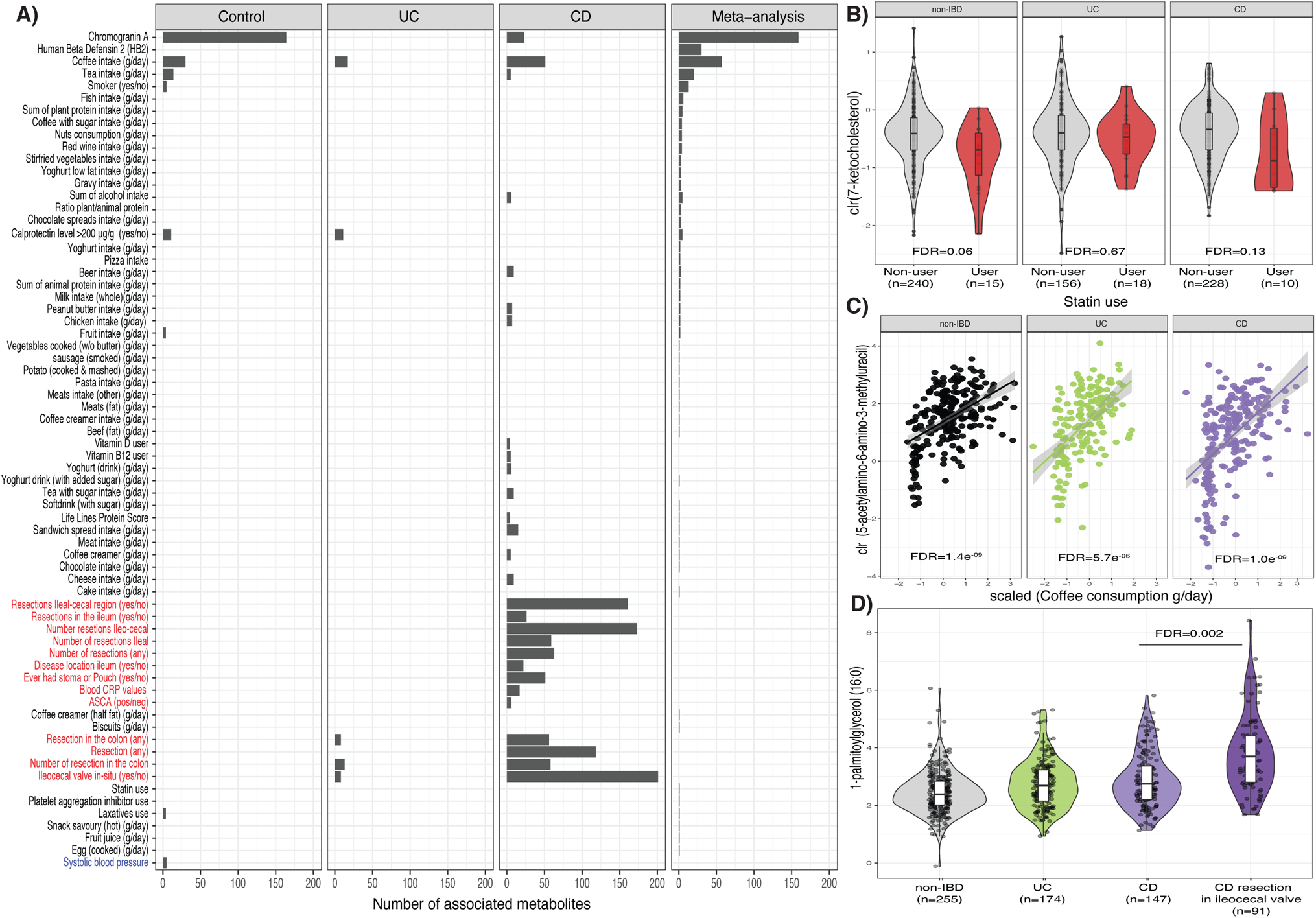
Potential determinants of faecal metabolite levels. **A.** Bar plot showing the number of significant associations between phenotypes and metabolites in each of the cohorts and in the meta-analysis (Suppl. Table 12. Only phenotypes with more than three associations are shown. Red labels indicate phenotypes exclusively available for cases and blue labels for controls. **B.** Boxplots depicting the levels of 7-ketocholesterol (expressed as clr-transformed AUC values) per cohort stratified by statin use. **C.** Correlation plot showing the relation between AAMU (expressed as clr-transformed AUC values) and coffee consumption (x-axis) per cohort. Coffee consumption is represented as the estimated consumption per day (grams/day) adjusted by overall individual calorie intake (see Methods). **D.** Boxplots showing the levels of 1-palmitoylglycerol (16:0). Boxplot shows the median and interquartile range (25^th^ and 75^th^). Whiskers show the 1.5*IQR range. Data distribution is represented by background violin-plot. Lines in the correlation plot show linear regression and shadows indicate the 95% confidence interval.

We also observed consistency between drug usage and drug-derived metabolites (**Suppl. Table 10**). O-desmethyltramadol, the main active metabolite of the opioid tramadol, was detected in several patients with CD using opioids (logistic regression, FDR = 0.009, **Suppl. Table 11**). In patients with UC, the use of mesalazine (5-aminosalicylate) was associated with higher levels of gentisate and N-acetylmethionine sulfoxide (linear regression, FDR = 0.005, **Suppl. Table 12**). Interestingly, 5-aminosalicylate has been reported as an alternative substrate of gentisate 1,2-dioxygenase, an enzyme involved in gentisate metabolism. Taken together, these results suggest that gentisate metabolism is significantly reduced in the presence of 5-aminosalicylate (**Suppl. Figure 3A**)^20^.

We found 158 associations between metabolites and dietary patterns (linear regression, FDR_meta_ < 0.05, **Suppl. Table 12**). Interestingly, approximately one-third of these were related to the consumption of coffee (n = 57), including positive correlations between coffee intake and the levels of picolinate and 5-Acetylamino-6-amino-3-methyluracil (AAMU), one of the major caffeine metabolites (**Fig 2C**) (linear regression, FDR_Meta_ < 0.05, **Suppl. Table 12**).

Beyond diet and medication, we detected cotinine in faeces of self-reported smokers (logistic regression, FDR_meta_ = 1.31e^−11^, **Suppl. Table 11**) and found that higher systolic blood pressure was associated with five molecules in controls, including higher agmatine levels (linear regression, FDR = 0.04, **Suppl. Table 12**).

### Intestinal resections are associated with long-term metabolic alterations

Although none of the patients in this cohort had had a recent surgical procedure at time of inclusion (average time between surgery and sample collection = 90 months, s.d. 87.4), having a history of intestinal resection(s) was strongly associated with faecal metabolite composition. In patients with CD, resection of the ileocecal valve was associated with changes in the abundance of 201 metabolites, including cholic acid and several monoacylglycerols. Colonic resection was also associated with modifications in the levels 56 molecules in CD and 8 molecules in UC (**Suppl. Table 12**, **Figure 2A,D**). For example, colonic resection negatively correlated with the faecal levels of pyridoxamine (vitamin B6) (**Suppl. Figure 3B**).

There were no significant differences in metabolites between different groups of disease behaviour or disease severity after statistically adjusting for gut surgery (resected vs non-resected). However, we did observe several interesting trends (linear regression, p-value < 0.05, FDR > 0.05, **Suppl. Table 13**). For example, patients with CD and penetrating diseases had lower butyrate levels (B1 vs B3). Disease severity (Montreal S score) also positively correlated with tyramine faecal abundance (**Suppl. Figure 3C**). In patients with UC, the extent of the disease (Montreal E score) was associated with the levels of four metabolites. For example, participants with proctitis (distal inflammation, E1 classification) had lower levels of 2R-3R-hydroxybutyrate and higher levels of cytidine compared to patients with extensive inflammation in the colon (proximal inflammation or pancolitis, E3 classification) (**Suppl. Figure 3D**). In addition, 33 metabolites were altered in patients with exclusively-colonic CD as compared with UC.

### *NAT2* genotype strongly associated with coffee metabolism

Next, we investigated the effect of host genetics on faecal metabolites by carrying out a faecal metabolome genome-wide association analysis. At a study-wide significance level (p-value < 3.12e^−10^), we found only an association between a genetic polymorphism located closely to *NAT2* (rs4921913) and AAMU (p-value_meta_ = 1.79e^−11^). This genetic variant is in linkage disequilibrium with a SNP reported to be associated with the ratio between 1,3-dimethylurate and AAMU (rs35246381, r^2^ > 0.8)^21^. As expected, we could also replicate this finding in our cohort (p-value_IBD_ = 8.46e^−09^, p-value_controls_ = 4.17e^−09^, p-value_meta_ = 3.57e^−13^, **Fig 3**). AAMU is a metabolite derived from coffee, and its levels in faeces are strongly correlated with coffee consumption. Nonetheless, this gene–metabolite association remained significant even after adjusting for coffee intake (p-value_IBD_ = 2.2e^−16^, p-value_controls_ = 2.0e^−09^).

**Figure 3.**
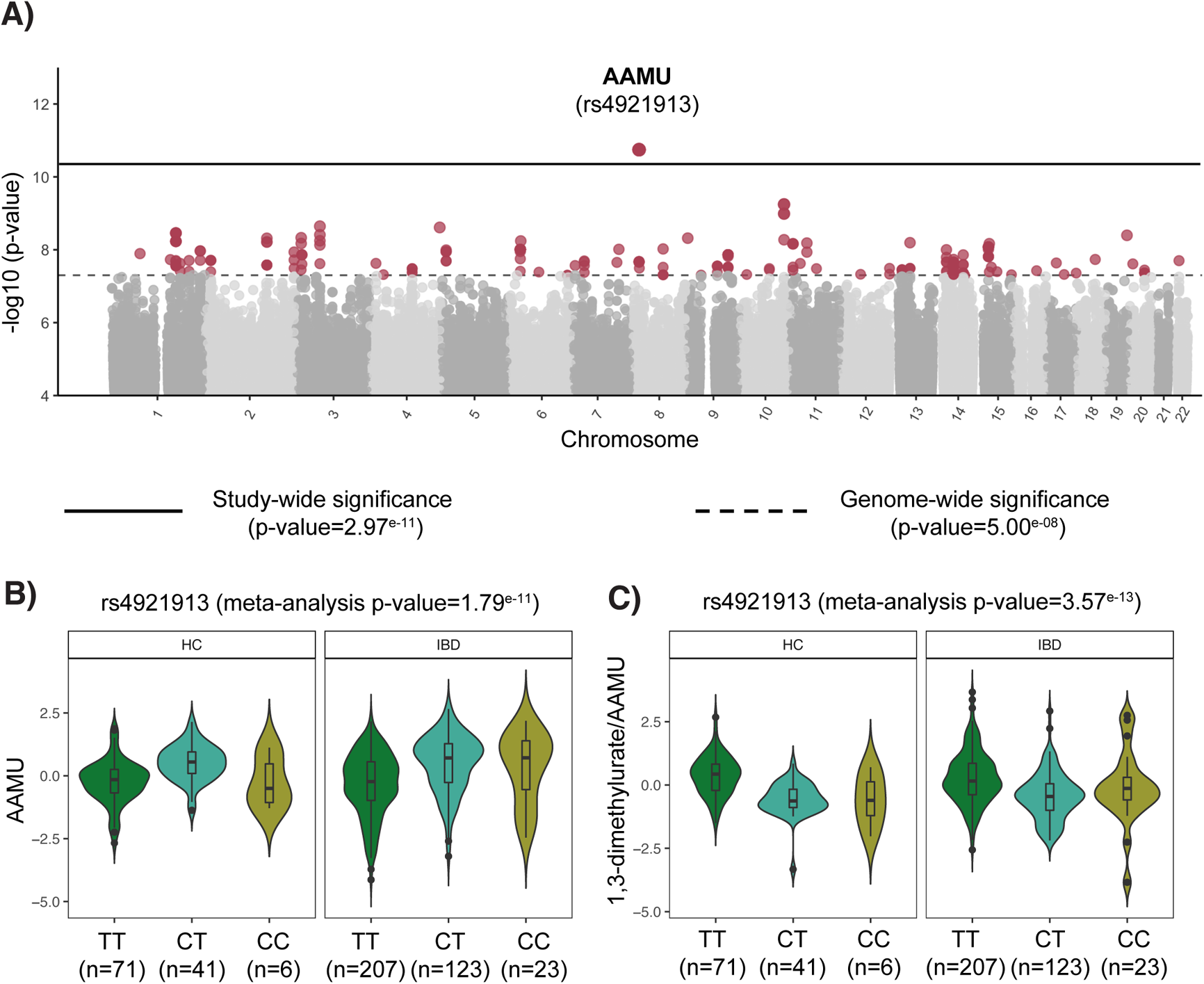
Genome-wide association between genetic polymorphisms and faecal metabolites. **A.** Manhattan plot shows the strong association between a single nucleotide polymorphism located in the *NAT2* gene and AAMU, a metabolite derived from caffeine. Solid horizontal line signifies the significance threshold corrected by multiple hypothesis testing. Dashed line indicates the classic genome-wide significance threshold. Metabolites passing this threshold (in red) are considered suggestive associations (Suppl. Table 14). **B.** Boxplot depicting the levels of AAMU in non-IBD controls and IBD, stratified by SNP rs4921913 genotype. **C.** Boxplot showing the relation between SNP rs4921913 and the ratio of 1,3-dimethylurate to AAMU. This association was previously described in the TwinsUK cohort^21^. Metabolite values are presented as the residuals of the model regressing the covariates age, sex, BMI and technical confounders. Boxplot shows the median and interquartile range (25^th^ and 75^th^). Whiskers show the 1.5*IQR range. Data distribution is represented by background violin-plot

At genome-wide significance level p-value < 5e^−08^, 267 genomic variants were associated with 61 different metabolites. For example, SNP rs4751995 in gene *PNLIPRP2* was associated with the level of a faecal choline derivative (1-palmitoyl-2-linoleoyl-digalactosyl glycerol (16:0/18:2), p-value_meta_ = 5.68e^−10^), and a genetic variant rs2331038 in gene *LRP5L* was associated with serotonin (**Suppl. Table 14**).

### Gut microbiota composition is linked to metabolomic profiles

We used neural networks to systematically assess the probability of co-occurrence between metabolites and bacteria. Higher cholic acid and creatinine levels correlated with an increase in the abundance of several pathobionts, including *Ruminococcus gnavus, Streptococcus mutans* and *Veillonella parvula*, and a decrease in *Methanobrevibacter smithii* and *Coprococcus eutactus* (**Fig 4A, Suppl. Table 15**).

**Figure 4.**
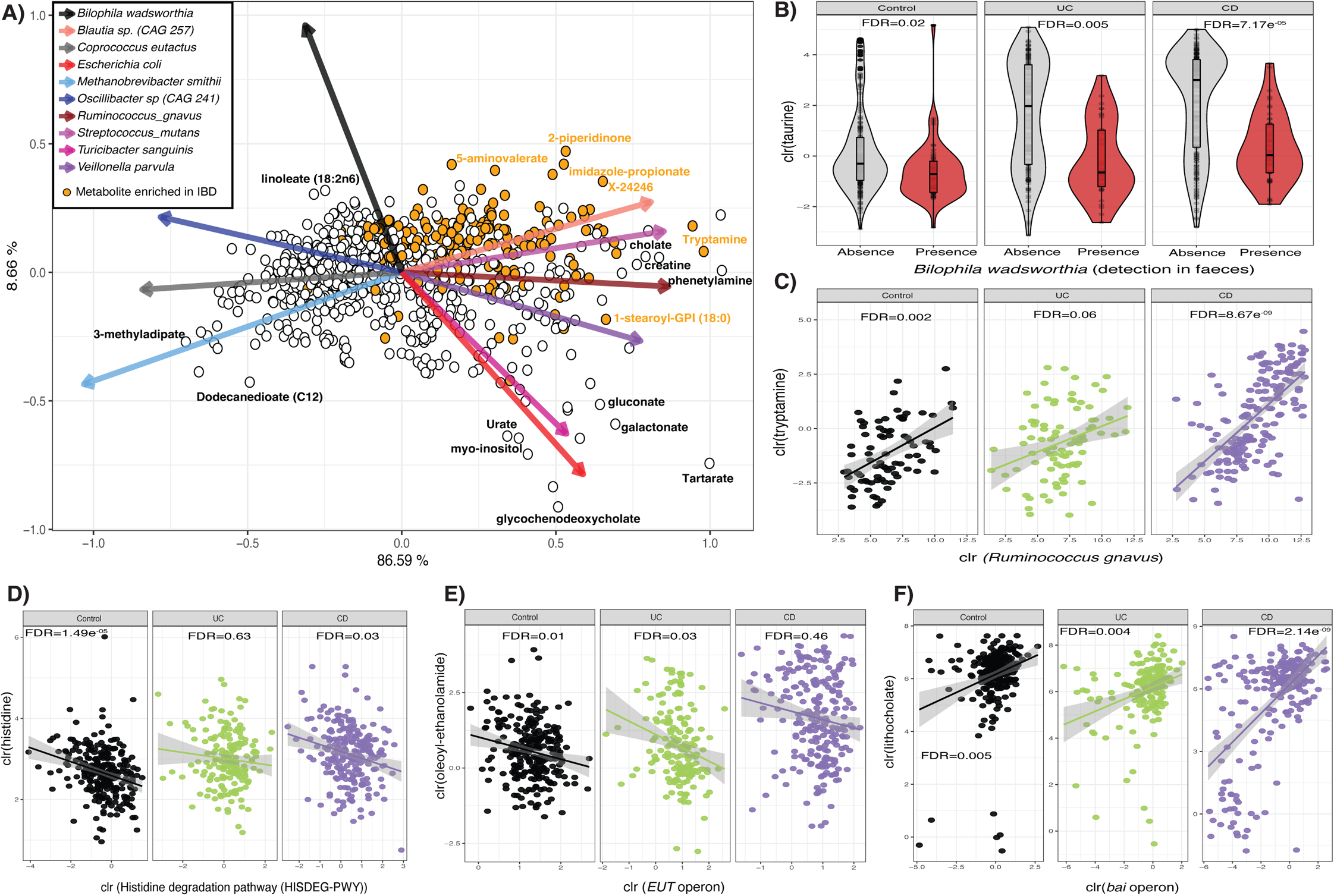
Metabolite co-occurrence with faecal microbes. **A.** Biplot representing conditional probabilities of co-occurrence between metabolites (dots) and microbes (arrows). Distances between dots and arrow tips represent the probability of co-occurrence of each metabolite and microbe. Orange dots highlight metabolites enriched in samples from patients with IBD in the linear regression analysis (see Methods, Suppl. Table 15). Arrow direction indicates the probability of microbes co-occurring with the levels of metabolites. To enhance interpretability, names of only a few metabolites are shown and only the top-10 species explaining the largest amount of variation are visualised. **B.** Taurine levels per cohort stratified by the presence or absence of *B. wadsworthia* in faecal metagenomes. **C.** Correlation between levels of tryptamine and abundance of *R. gnavus*. Only samples in which the bacterium had a non-zero relative abundance are shown. **D-F.** The relation between histidine and MetaCyc Histidine degradation pathway (**D**), between oleoyl-ethanolamide and the *eut* operon (**E**) and between lithocholic acid and the *bai* operon (**F**) are shown as examples of the correlation between microbiota metabolic potential and metabolite levels, per cohort. Metabolite, bacteria and pathway values are clr-transformed. Boxplot shows the median and interquartile range (25^th^ and 75^th^). Whiskers show the 1.5*IQR range. Data distribution is represented by background violin-plot. Correlation plot lines show linear regression. Shadows indicate the 95% confidence interval.

We then tested if the microbiome–metabolite co-occurrences were consistently observed in the three conditions in our study (CD, UC and non-IBD controls). First, we tested if the presence/absence of specific taxa in faeces was associated with the levels of faecal metabolites. The meta-analysis revealed 16,237 associations (**Suppl. Figure 5**), of which 689 were significant in all three conditions (FDR_controls_ < 0.05, FDR_CD_ < 0.05, FDR_UC_ < 0.05, **Suppl. Table 16**). The presence of *Akkermansia municiphila* and *Oscillibacter sp*. *(CAG 241)* were associated with higher levels of fatty acids (fatty acyls: 3-methyl adipate, azelate (C9-DC), sebacate (C10-DC) and dodecanedioate (C12)). The presence of *Bilophila wadsworthia* was associated with lower levels of taurine, betaine and N,N,N-trimethyl-L-alanyl-L-proline (TMAP) (**Fig 4B**, linear regression, FDR_meta_ < 0.05).

Additionally, we correlated non-zero microbial abundances with levels of metabolites and found 2,303 significant positive associations (**Suppl. Table 17**, FDR < 0.05). For example, the abundances of *Blautia wexlerae* and *Dorea formicigenerans* were positively correlated with lysine degradation products (N6-carboxymethyl lysine and N6,N6-dimethyl lysine), while *F. prausnitzii* correlated with hypoxanthine levels (linear regression, FDR_meta_ = 0.006). The levels of tryptamine were positively correlated with the abundance of *R. gnavus* (linear regression, FDR_meta_ = 4.87e^−08^) (**Fig 4C**) and levels of imidazole propionate with the abundance of *Streptococcus parasanguinis* (linear regression, FDR_meta_ = 2.86e^−04^).

When exploring the metabolic potential of the gut microbiota, we observed significant differences in the abundances of 93 pathways and 94 gene clusters between cases and controls (**Suppl. Table 18**). In IBD, these alterations were characterised by increased amino acid utilisation and decreased fermentation pathways and fatty acid metabolism. For example, histidine degradation pathways (MetaCyC ID: HISDEG and PWY-5030) and histidine utilisation operon (hutHGIU) were enriched in both UC and CD metagenomes. The increase of the L-histidine degradation pathway I (HISDEG) was negatively correlated with the levels of histidine (linear regression, FDR_meta_ = 4.0e^−04^), although it was not significantly associated with the levels of imidazole propionate (**Fig 4C, Suppl. Table 19**). In addition, the abundances of ethanolamides were negatively correlated with the ethanolamine utilisation operon (eut). Ethanolamines are components of host cell membranes, and their levels increase during periods of inflammation. The *eut* operon is known to be carried by several gut pathobionts, allowing the use of ethanolamine as a source of carbon and nitrogen, which might provide a selective advantage during periods of active disease (**Fig 4D**)^22^. The abundance of *bai* operons was associated with higher levels of faecal lithocholic acid (linear regression, FDR_meta_ = 0.003) and 12-ketolithocholate (linear regression, FDR_meta_ = 0.005) and lower levels of cholic acid (linear regression, FDR_meta_ = 3.54e^−05^) (**Fig 4E**) (**Suppl. Table 20**).

### Metabolic alterations in dysbiosis

Considering that the gut microbiota of patients with IBD undergo transitions from eubiosis to dysbiosis, which might precede or indicate periods of active disease^3,23^, we investigated metabolite differences between patients with dysbiotic and eubiotic microbial compositions. In total, we identified 191 participants with dysbiotic microbiota (**Fig 5A,B**). The clinical characteristics of these participants showed an enrichment of CD subtype (n = 130, chi-squared test FDR = 2.45e^−04^) and ileocecal valve resections (n = 76, chi-squared test, FDR = 2.93e^−10^). Levels of chromogranin A were also higher in dysbiotic patients (Wilcoxon-test, FDR = 4.96e^−05^). However, no significant differences were observed between groups in the levels of faecal calprotectin (chi-squared test FDR = 0.65, **Suppl. Table 5**).

**Figure 5.**
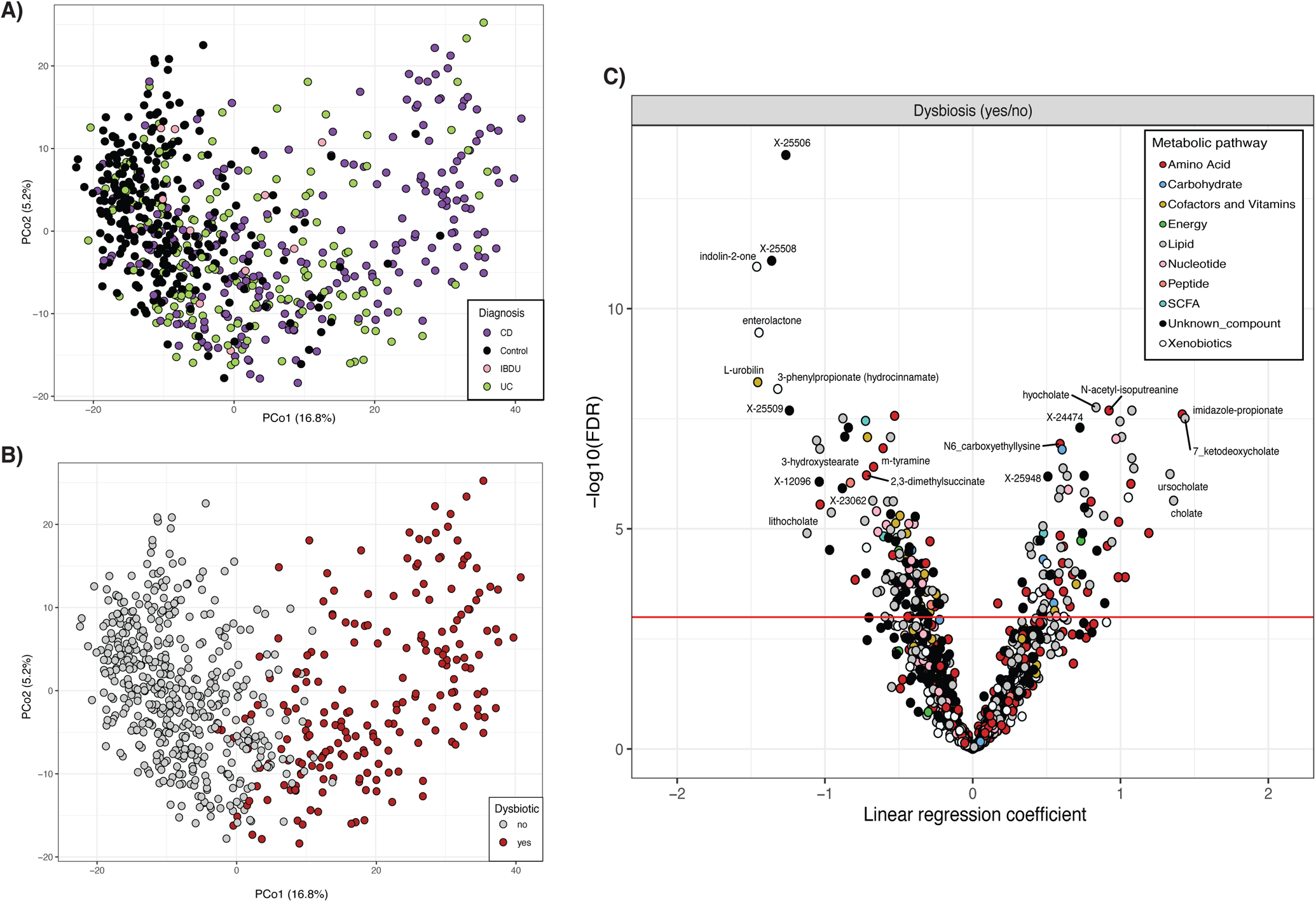
Metabolic signature of patients with intestinal dysbiosis. **A.** Principal coordinate analysis on microbiome composition per sample (dots). Colours indicate disease phenotypes: CD (purple), UC (green), IBD-undetermined (pink), non-IBD (black). **B.** Red dots depict samples considered to be dysbiotic based on the median distance to non-IBD samples. **C.** Volcano plot showing the p-value (y-axis) and regression coefficients (x-axis) of the association analyses between dysbiotic and non-dysbiotic IBD samples (Suppl. Table 5). Dot colour indicates pathway annotations provided by Metabolon.

**Figure 6.**
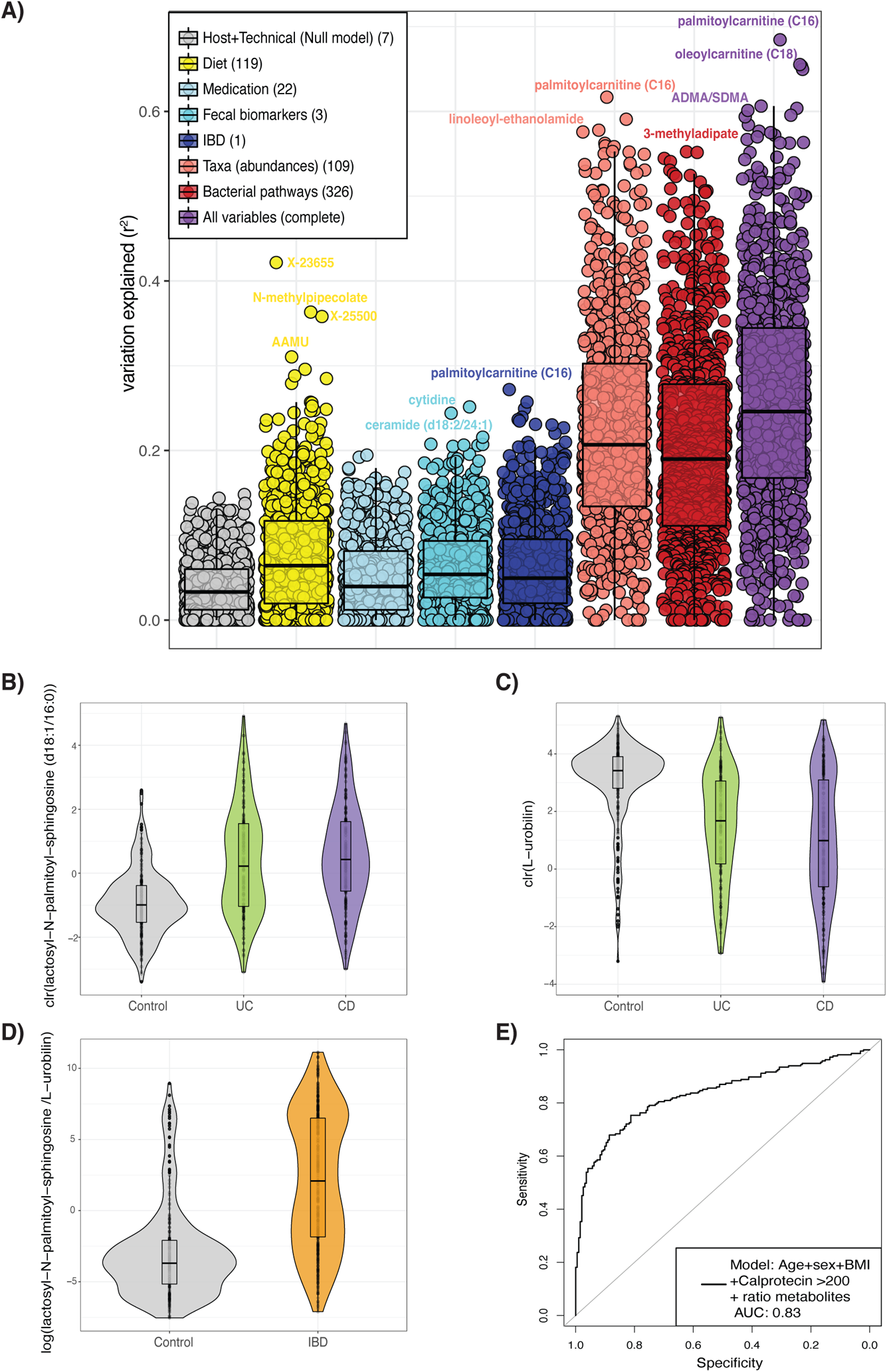
Metabolite prediction and biomarker discovery for the diagnosis of IBD. **A.** Microbial abundances (light red) and bacterial pathways (dark red) show the largest capacity to predict the levels of metabolites. Boxplots show the capacity to predict metabolites levels of eight different models using seven types of data. Dots represent metabolites, and values in the y-axis represent the percentage of variation explained from cross-validated penalised regression methods using different sets of predictors (see Methods). The number of features in each model are indicated in parentheses in the legend (Suppl. Table 21). **B-C**. Show the abundance of the metabolites with the highest potential to discriminate between samples from non-IBD (grey) and IBD (UC in green and CD in purple). **D.** Boxplots depict the value of a potential biomarker for IBD. Y-axis is the log-transformed value of the ratio constructed from the levels of lactosyl-N-palmitoyl-sphingosine (d18:1/16:0) and L-urobilin. Boxplot in grey depicts values in non-IBD controls. Boxplot in orange depicts values in patients with IBD. **E.** Receiver operating characteristic curve (ROC curve) of the prediction model based on patient characteristics (age, sex and BMI), the levels of faecal calprotectin (expressed as a binary trait (yes/no) if levels of this marker were > 200 μg/g of faeces) and the ratio between metabolites. The prediction value, expressed as the area under the curve (AUC), reached a value of 0.83 in the test dataset. Metabolite values are clr-transformed. Boxplot shows the median and interquartile range (25^th^ and 75^th^). Whiskers show the 1.5*IQR range. Data distribution is represented by background violin-plot.

After correcting for disease phenotype (CD or UC), history of surgery, integrity of the ileocecal valve and dietary differences between the two groups, we identified 202 enriched metabolites and 258 depleted metabolites in patients with dysbiosis. In patients with dysbiosis, levels of SCFA (hexanoic, valeric and butyric acid), indolin-2-one and 3-phenylpropionate (hydrocinnamic acid) were reduced and N-acetyl-isoputreanine, long-chain polyunsaturated fatty acids and bile acids were significantly increased (linear regression, FDR < 0.05, **Suppl. Table 5**). In addition, this group of patients showed a higher prevalence of taurine-conjugated and sulphated bile acids (logistic regression, FDR < 0.05, **Suppl. Table 5**). Overall, we observed lower levels of secondary bile acids compared to primary bile acids in dysbiotic samples, suggesting an accumulation of primary bile acids during periods of dysbiosis (linear regression, FDR_DCA:CA_ = 0.007, FDR_LCA:CDCA_ = 9.74e^−04^).

### Microbiome composition predicts metabolite levels in faeces

In light of the numerous associations between metabolite levels and microbes, we explored to what extent metabolites could be predicted from metagenomic datasets. Age, sex, BMI, average bowel movements a day and technical experimental factors explained 3% of the metabolite variation. Dietary habits could predict > 20% of the variation in cysteine (25%, s.d. 1%), bilirubin (21%, s.d. 8.1%) and metabolites related to coffee consumption, including an unidentified metabolite that correlates with N-methyl pipecolic acid (X-23655, 42%, s.d. 6%) and AAMU (31%, s.d. 9%). IBD (expressed as a binary trait) explained > 20% of the variation in palmitoylcarnitine (C16) and 15 other metabolites (**Suppl. Table 21**). In contrast, bacterial abundances were a strong predictor of 82 metabolites (> 40% of the variation), including of the levels of molecules such as lithocholate (41%, s.d. 18%) and dimethylarginine (ADMA/SDMA, 53%, s.d. 4%). Adding diet and participant characteristics to the model provided only a slight increase in the prediction power compared to that of bacterial abundance alone (paired Wilcoxon-test, p-value < 2.2×10^−6^) (**Fig 5A**).

### Faecal metabolomic profiles correctly classify IBD samples

To discover potential biomarkers, we used a machine learning approach to identify metabolites that could predict disease phenotypes (see Suppl. Methods). Including the ratio between the sphingolipid lactosyl-N-palmitoyl-sphingosine (d18:1/16:0) and L-urobilin improved the accuracy of age, sex, BMI and faecal calprotectin levels as disease predictors (AUC_cv_ = 0.85, AUC_test_ = 0.83, p-value = 9.89e^−13^, **Fig 5 B-E, Suppl. Table 22**), and a similar performance was achieved with bacteria abundances (AUC_cv_ = 0.86, AUC_test_ = 0.84, p-value = 6.04e^−14^, **Suppl. Figure 4 B,C**). A combination of three species—*Flavonifractor plautii*, *Gemmiger formicilis* and *Methanobrevibacter smithii*—was identified as the most discriminative feature between cases and controls. Combining metabolite and microbiome ratios led to a modest but significant increase in model performance (AUC_test_ = 0.85, p-value = 4.34 e^−09^).

Within patients with IBD, metabolites showed a limited power to correctly classify CD or UC samples (AUC_cv_ = 0.78, AUC_test_ = 0.67) and active disease vs remission (AUC_cv_ = 0.72, AUC_test_ = 0.60) (**Suppl. Table 22**).

## DISCUSSION

We comprehensively characterised faecal metabolites in samples from patients with IBD and representatives of the Dutch population. Our results revealed alterations in the levels of more than 300 highly prevalent faecal metabolites in patients with IBD. Additionally, we described potential determinants of faecal metabolome composition by integrating untargeted metabolomics with extensive information on dietary habits, host genetics, clinical characteristics and gut microbiota composition.

The drastic alteration in faecal metabolite composition in patients with IBD suggests a switch from a saccharolytic to a proteolytic fermentation metabolism^24^. We observed increased levels of products of amino acid metabolism. For example, patients with IBD showed higher levels of p-cresol sulphate compared to controls (FDR_CD_ = 1.45e^−04^, FDR_UC_ = 1.54e^−03^, **Suppl. Table 7**). This compound originates from fermentation of aromatic amino acids in the large intestine^25^, and its accumulation has been related to several diseases, including chronic kidney disease^26^ and colorectal cancer^27,28^. Phenol sulphate, a uremic toxin derived from metabolism of tyrosine^29,30^, was more prevalent in faeces of patients with CD (logistic regression, FDR_CD_ = 0.003, **Suppl. Table 7**). Other metabolites enriched in the faeces of patients with IBD included molecules with a known pro-inflammatory effect, such as histidine, polyunsaturated fatty acids and sphingosines. In agreement with previous reports, we also found a strong enrichment of ceramides, polyamines, acylcarnitines and ethanolamides in the faeces of patients with CD and depletion of SCFA and gamma-glutamyl amino acids in UC (**Suppl. Table 7**, FDR<0.05)^3–5,7,31,32^.

The strong correlation between microbial composition and faecal metabolites suggests that metabolomic data partially capture bacterial activity in the gut. We found *B.wadsworthia* to be associated with taurine, betaine and TMAP levels (**Suppl. Table 16**). While this bacterium has the capacity to metabolise taurine and betaine, little is known about TMAP^33^. It has recently been shown that *B. wadsworthia* can convert trimethylamine into dimethylamine and consequently reduce the overall production of trimethylamine-N-oxide (TMAO)^34^. Therefore, we hypothesise that the demethylation capacities of this bacterium might explain the association between TMAP and *B. wadsworthia*.

Another example of the strong correlation between metabolites and bacteria is the positive correlation we observe between levels of hypoxanthine, butyric and acetic acids, and the abundance of *F. prausnitzii* (**Suppl. Table 17**). Hypoxanthine is a molecule that contributes to maintaining the intestinal epithelium^35^. Hypoxanthine can be produced by *F. prausnitzii* through the metabolization of adenine^36^. Moreover, the relative abundance of *Ruminococcus gnavus* was related to tryptamine levels (**Suppl. Table 17**). *R. gnavus,* a bacterium highly abundant in dysbiotic periods in patients with IBD^37^, can produce tryptamine by decarboxylation of tryptophan^38^. Accumulation of tryptamine can increase gut motility via activation of serotonin receptor-4, which may explain why some patients experience decreased intestinal transit times during flares^39^. Additionally, the positive correlation we observe between *S. parasanguinis* and imidazole propionate could be explained by the capacity of this bacterium to degrade histidine^40^. Imidazole propionate has been associated with the risk of developing type 2 diabetes and regulates activation of the mTORC1 signalling pathway^41,42^, which is implicated in IBD^43^.

The relation between the microbiome and metabolites allowed us to predict the levels of 82 faecal metabolites using metagenomic sequencing data (**Suppl. Table 21**). Consistent with observations in other cohorts^21,44–46^, well-predicted molecules included putrescine, urobilin, bile salts and fatty acids. However, and in agreement with Muller et al.^46^, models trained in controls and tested in IBD showed lower prediction accuracy than models trained with both IBD and non-IBD datasets. The low cross-predictability between cases and controls implies that some microbiota– metabolite relations might be context-specific. In fact, when clustering samples either based on microbiome or metabolomic profiles, IBD samples tend to constitute two clusters—one overlapping with controls and another almost exclusively populated by samples of patients with IBD or dysbiosis (**Fig 1A, Fig 5A**). This pattern was also observed in the study of Jacobs et al.^47^ on faecal samples from patients with IBD and their unaffected first-degree relatives. Interestingly, in our cohort, the dysbiotic cluster was mainly populated by samples from patients with ileum disease involvement and/or an ileocecal valve resection (**Suppl. Table 5**). Accumulating literature demonstrates that disruptions in the small intestine due to inflammation or surgery impact the metabolite and microbial composition in faeces^48,49^. Halfvarson and colleagues^23^ showed that patients with intestinal surgery in the ileum had a less stable microbiota and more often transited between non-IBD and IBD-enriched clusters. We observed that the dysbiotic cluster presented an enrichment of primary bile acids as compared to eubiotic samples from patients with IBD (**Suppl. Table 5**). It has been reported that an increase in cholic acid can exert selective pressure on the gut ecosystem due to its antimicrobial properties^50^. Overall, and considering the crucial role of the small intestine in nutrient absorption, we hypothesise that perturbations in this section of the gut lead to long-lasting alterations in the concentrations of bile acids, amino acids and lipids in the colon, which might re-shape the microbial composition towards a dysbiotic state.

Along with the influence of the gut microbiota, diet and lifestyle are potential determinants of the abundance of small molecules in the human body. By correlating metabolites to dietary data, medication use and lifestyle factors, we found that daily habits such as smoking or coffee and tea consumption strongly correlated with their derivative molecules (**Suppl. Table 12**). Moreover, N-acetyl proline levels were associated with wine consumption. N-acetyl proline is an alpha-amino acid product of proline which is highly abundant in grape juice and wine^51^. In addition, dipeptides were more abundant in participants consuming more meat. Anserine, for example, which is present in the muscle tissue of poultry^52^, was found in higher levels in participants who consume chicken (**Suppl. Table 12**). Despite these associations, long-term dietary habits show only a moderate association with faecal metabolome composition. We hypothesise that our dietary data underestimates the contribution of food intake to levels of faecal metabolites since it is based on food frequency questionnaires. Future studies should consider the use of 24-hour dietary recalls to capture daily dietary variations when aiming to explore relations between food intake and faecal metabolites and food–microbiome interactions.

Host genetics showed only a small impact on metabolite levels in faeces. The only association that passed our significance threshold was between a single nucleotide polymorphism near the *NAT2* gene and AAMU, a caffeine metabolism product (**Suppl. Table 14**). *NAT2* encodes an N-acetyltransferase enzyme that detoxifies several xenobiotics, including coffee and certain types of medication. A study in the TwinsUK biobank also reported this association and estimated that host genetics has a moderate effect on faecal metabolites, with an average heritability of ~18%^21^. This relatively low heritability contrasts with the impact of host genetics on the levels of circulating metabolites^44,53^ and might be explained by the fact that faecal metabolites are primarily influenced by microbial transformations occurring in the colon, which can potentially mask genetic effects. Moreover, sample sizes are still a limiting factor for discovering metabolite–genome associations. In fact, when using a looser significance cut-off (p-value < 5×10^−8^), we found > 200 suggestive associations pointing to the metabolism of cholesterol and serotonin. For example, *LRP5L* was associated with serotonin and *PNLIPRP2* with 1-palmitoyl-2-linoleoyl-digalactosyl glycerol (16:0/18:2) (**Suppl. Table 14**). *LRP5L* belongs to the LDL receptor family found to be involved in controlling serotonin levels in the duodenum^54^. Both *PNLIPRP2* and 1-palmitoyl-2-linoleoyl-digalactosyl glycerol (a choline derivative) are linked to cholesterol metabolism, supporting the fact that choline supplements maintain blood cholesterol homeostasis^55^, and PNLIPRP2 has been associated with LDL levels in blood^56^.

Finally, considering that colonoscopies are still the gold standard for diagnosing IBD, we demonstrated the potential of faecal metabolites as a non-invasive method for disease diagnosis. The ratio between the levels of two metabolites, lactosyl-N-palmitoyl-sphingosine (d18:1/16:0) and L-urobilin or stercobilinogen was identified as a biomarker for IBD in our cohort (**Suppl. Table 22**). Reduced levels of L-urobilin and increased sphingolipids in faeces of patients with IBD have been observed in other cohorts^3,5^. Faecal measurements targeting these two commonly detected molecules could be relatively easy to implement in the clinic. These could, in combination with the faecal calprotectin levels, increase the accuracy of non-invasive tests.

In conclusion, this study provides a detailed characterisation of the faecal metabolites in the context of health and intestinal inflammation, replicating known disease-relevant molecules and expanding our knowledge of disease heterogeneity. In addition, we pinpoint multiple associations between host, microbiota, diet and faecal metabolite levels, which we believe provide valuable resources for further investigation of metabolite- or microbiota-based interventions and treatment in IBD.

## Authors contribution

AVV and RKW designed the study. AVV, SH, SAS, RG, HEA contributed to the data analysis. AVV and SH wrote the manuscript. BHJ handled the samples in the laboratory. LB, SH, SAS, VC, HEA, JF, AZ, AAG, JP, JS, CG, GA, RAAAR and RKW critically reviewed the manuscript.

## Ethical approval

All participants signed an informed consent form prior to sample collection. Institutional ethics review board (IRB) approval was available for Lifelines DEEP (ref. M12.113965) and 1000 IBD (IRB-number 2008.338) cohorts.

## Data availability

Tables containing the levels of faecal metabolites and bacterial taxa abundances are provided with the manuscript. The raw metagenomics, host genomics and phenotypic data used in this study are available from the European Genome–Phenome Archive data repository: 1000 Inflammatory bowel disease (IBD) cohort [https://www.ebi.ac.uk/ega/datasets/EGAD00001004194], Lifelines DEEP cohort [https://www.ebi.ac.uk/ega/datasets/EGAD00001001991]. Datasets are available upon request to the University Medical Center of Groningen (UMCG), LifeLines. This includes submitting a letter of intent to the corresponding data access committees.

Codes and processed microbiome and metabolomics data are publicly available at: https://github.com/GRONINGEN-MICROBIOME-CENTRE/Fecal_Metabolites_IBD

## SUPPLEMENTARY FIGURES

**Suppl. Figure 1.**
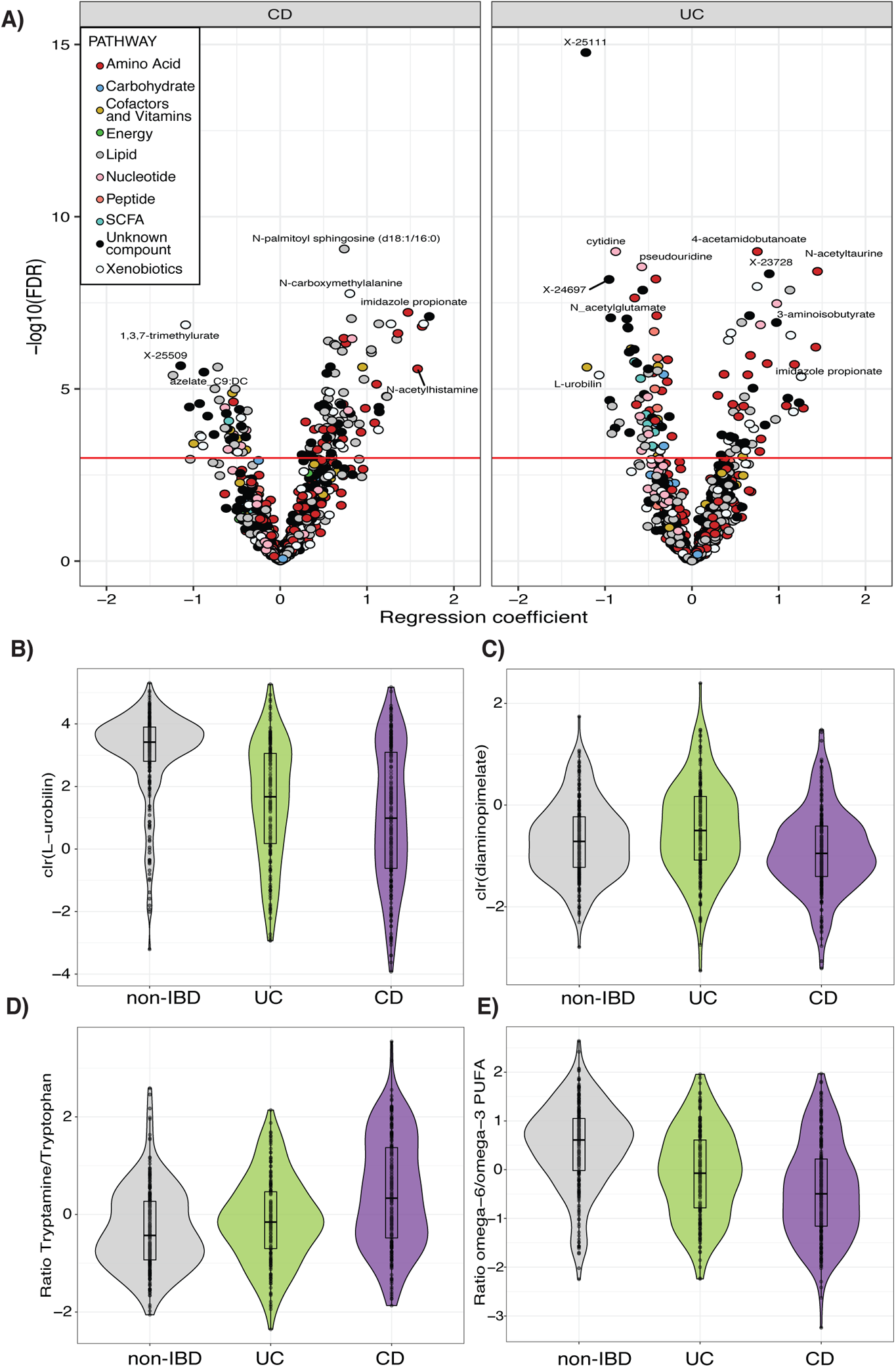
Metabolite alterations associated to IBD. **A.** Volcano plot showing the p-value (y-axis) and regression coefficients (x-axis) of the association analyses between cases (CD and UC) and non-IBD controls. Dot colour indicates the pathway annotation provided by Metabolon. **B-C.** Levels of L-urobilin and diaminopimelate stratified by disease phenotype. Metabolite levels are clr-transformed. L-urobilin is significantly decreased in both UC and CD and diaminopimelate significantly increased in UC. **D.** Log-ratio between tryptamine and tryptophan levels. This ratio was significantly increased in patients with CD. **E.** Patients with IBD show lower levels of the log-ratio between omega-6 and omega-3 polyunsaturated fatty acids.

**Suppl. Figure 2.**
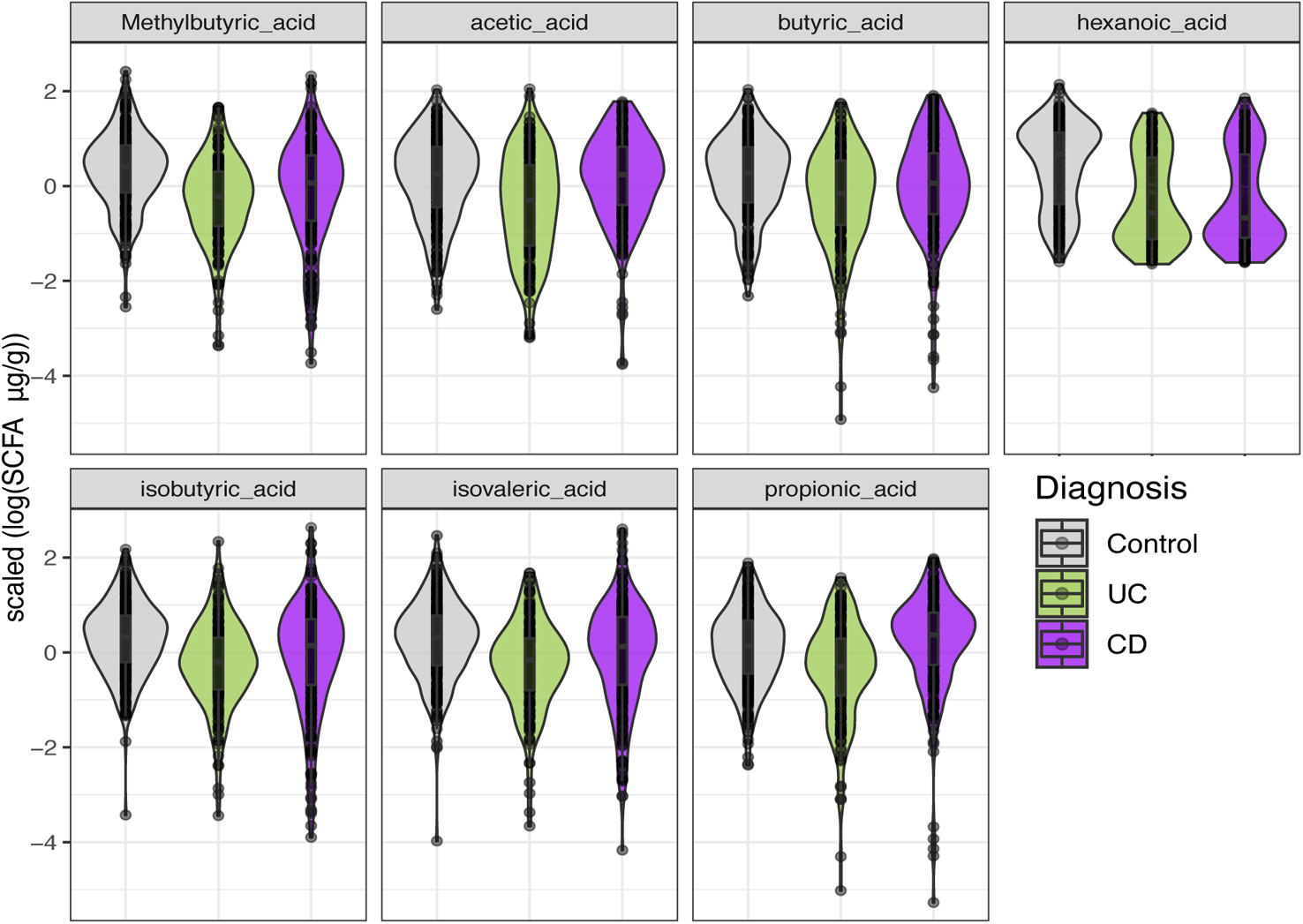
Short-chain fatty acids are depleted in patients with UC. Boxplots showing the concentrations of short-chain fatty acids per cohort. For visualisation purposes, metabolite concentrations are log-transformed and scaled.

**Suppl. Figure 3.**
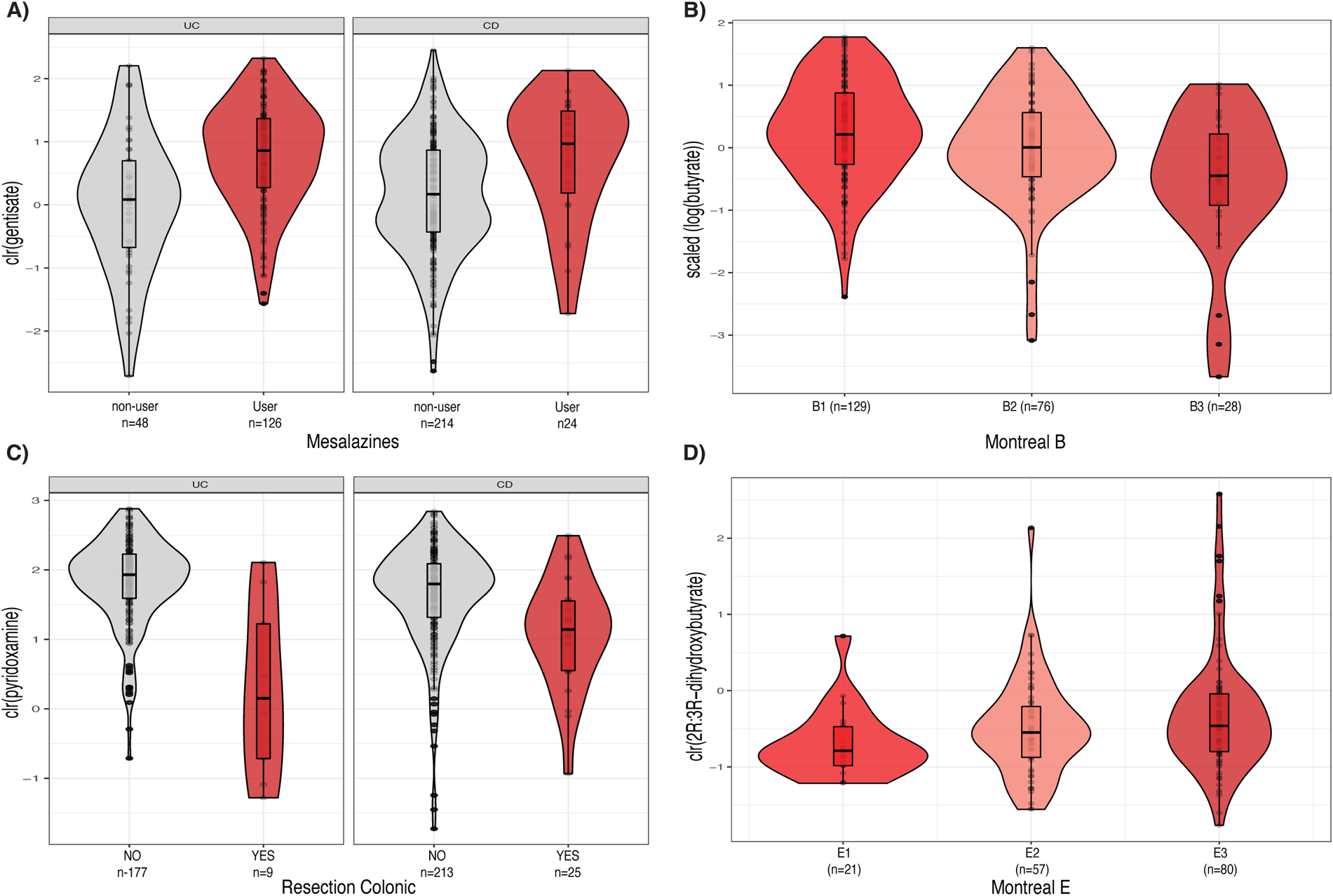
Metabolite levels stratified by disease and patient characteristics. **A.** Gentisate levels are increased in patients with IBD who used mesalamines. Y-axis represents clr-transformed metabolite levels. **B**. Levels of butyrate are lower in patients with CD and penetrating disease (Montreal B3) compared to non-structuring and non-penetrating forms of the disease (Montreal B1). **C**. Boxplots showing that patients with resections in the colon show lower vitamin B6 levels (pyridoxamine). **D**. In patients with UC, levels of 2R:3R-dihydroxybutyrate show a tendency to positively correlate with disease extension.

**Suppl. Figure 4.**
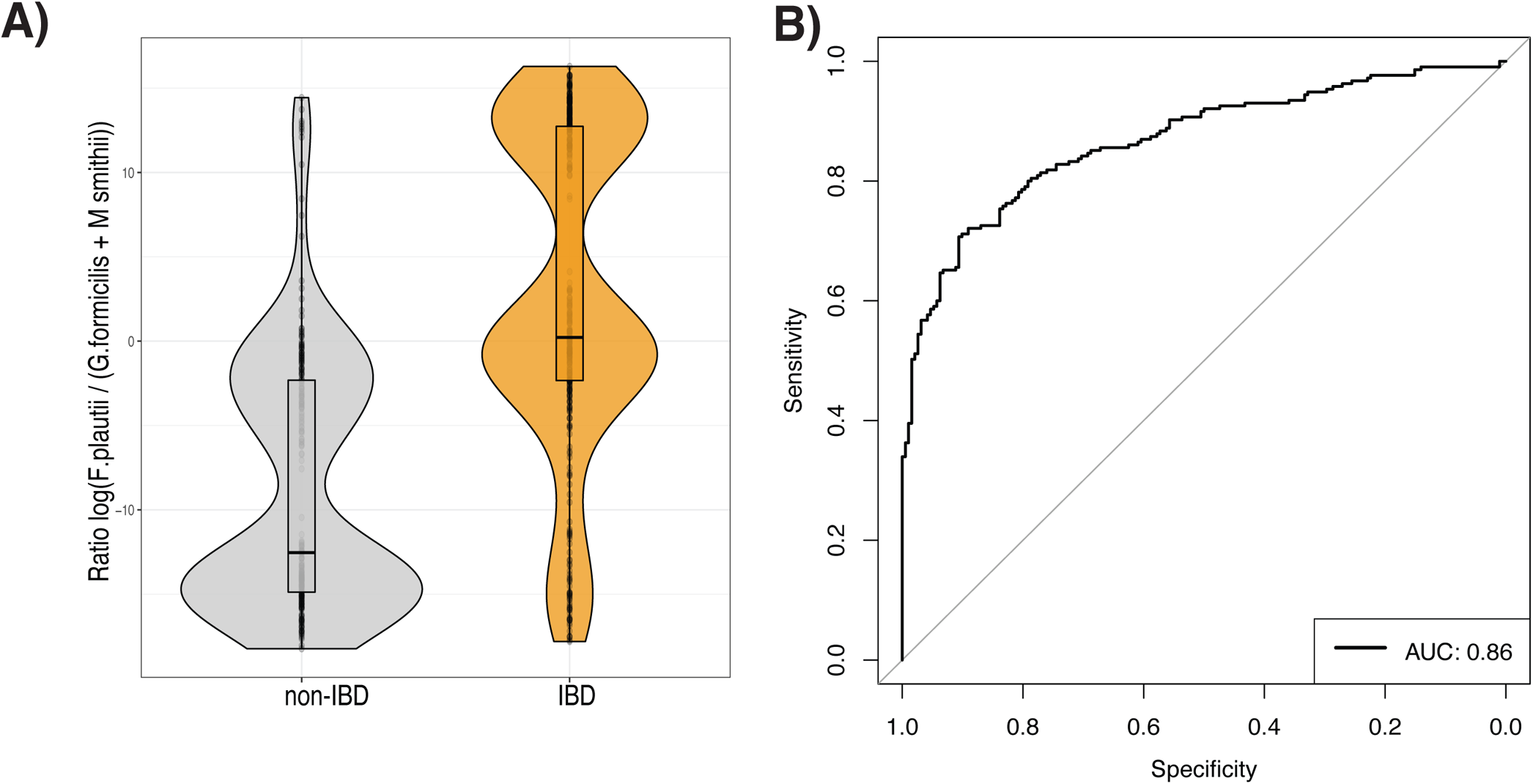
Bacterial taxa abundances can be used to classify non-IBD and IBD faecal samples. **A.** Boxplot depicts the values of a potential biomarker for IBD. Y-axis represents the log-transformed values of the ratio constructed from the abundance of *F. prausnitzii* and the sum of *G. formicilis* and *M. smithii*. Boxplot in grey shows values in non-IBD controls. Boxplot in orange shows values in samples from patients with IBD. **B.** Receiver operating characteristic curve (ROC curve) of the prediction model based on patient characteristics (age, sex, BMI), the levels of faecal calprotectin (expressed as a binary trait (yes/no) if levels of this marker were > 200 μg/g of faeces) and the ratio based on bacterial taxa. The prediction value, expressed as the area under the curve (AUC), reached a value of 0.86 in the test dataset. Metabolite values are clr-transformed. Boxplot shows the median and interquartile range (25^th^ and 75^th^). Whiskers show the 1.5*IQR range. Data distribution is represented by background violin-plot.

**Suppl. Figure 5.**
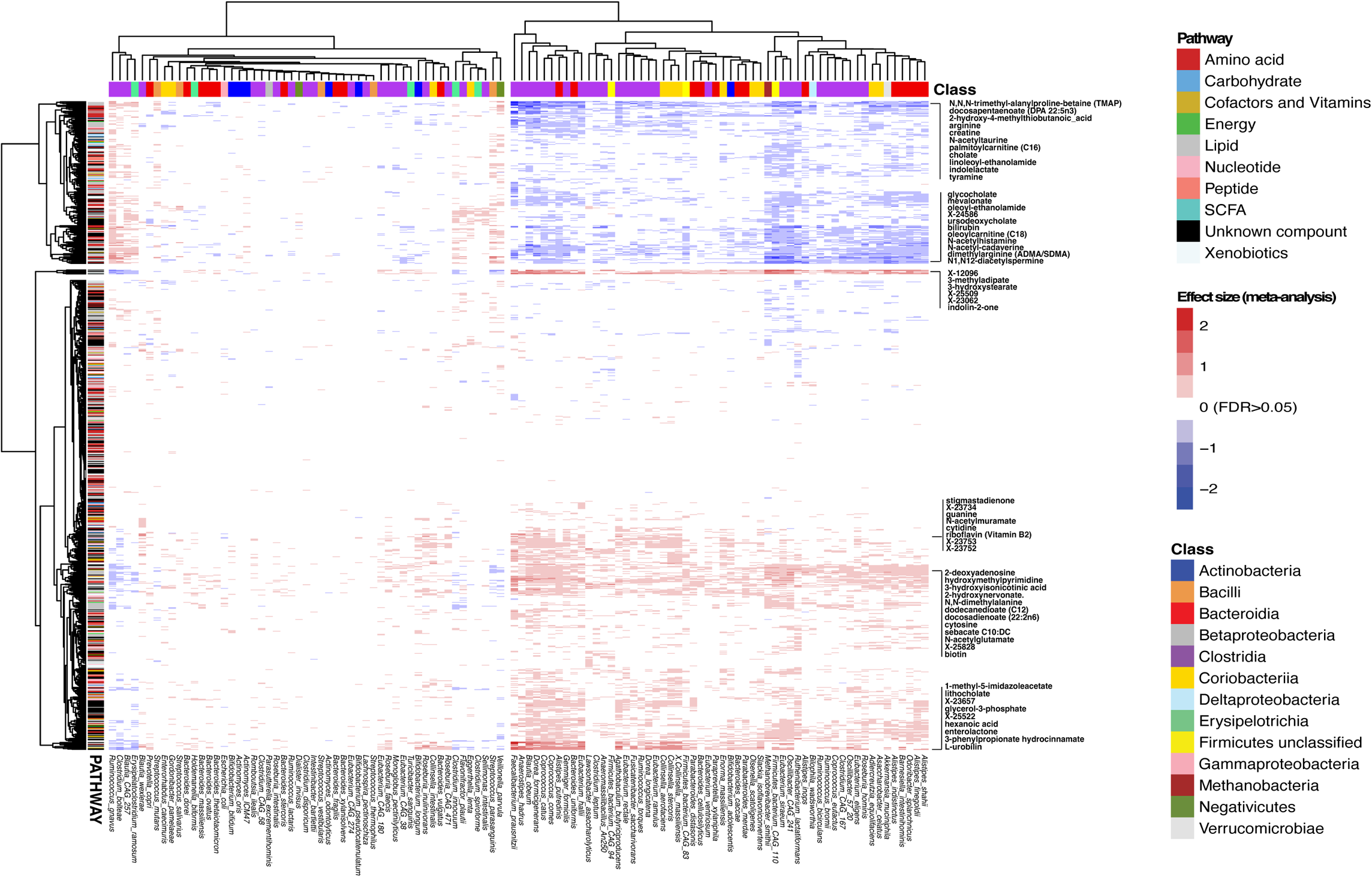
Heatmap showing co-occurrence of metabolites and bacteria in faeces. Heatmap showing two distinct patterns between the presence of bacteria and the levels of metabolites. Metabolites on the x-axis are annotated according to the pathway annotation provided by Metabolon. On the y-axis, bacteria are annotated based on their taxonomic class. Clustering is based on the regression coefficient derived from the meta-analysis of the relation between metabolite levels and detection or absence of a species in the faecal samples in each dataset (CD, UC, non-IBD). Red cells indicate positive associations and blue cells indicate negative associations between species and metabolites. White cells represent non-significant associations in the meta-analysis. For readability, only the labels of certain metabolites are shown.

**Suppl. Figure 6.**
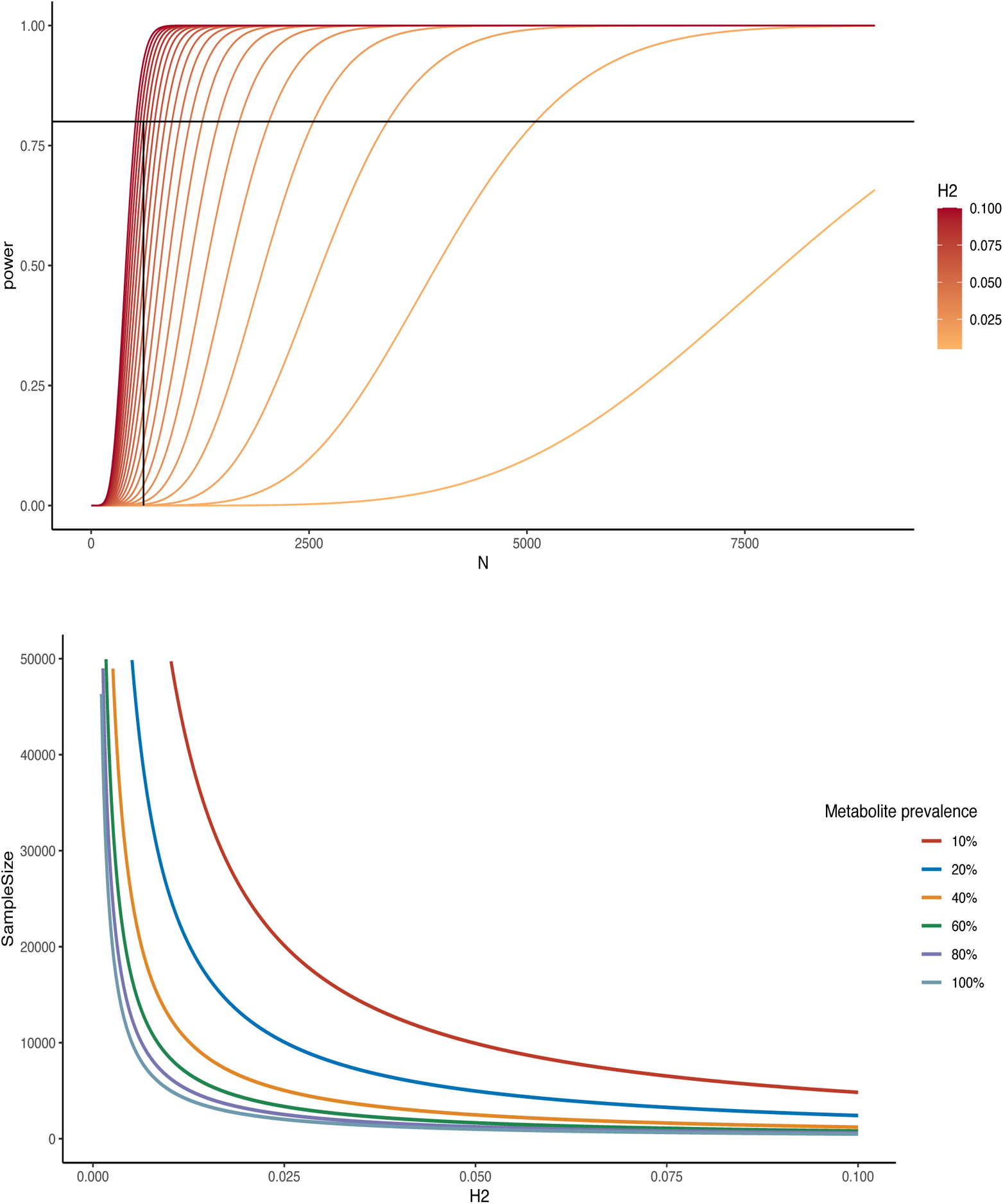
Statistical power to detect associations between host genetics and faecal metabolites. **A.** Power to detect associations dependent on sample size (alpha = 3.14e^−10^, variance explained by genetic effect (h^2^) = 0.08). **B.** Sample size to detect associations dependent on h^2^ considering different degrees of missingness from 10%–100% across all samples (alpha = 3.14e-10, power = 0.8).

## SUPPLEMENTARY TABLES

**Suppl. Table 1.** Cohort description and phenotypes

**Suppl. Table 2.** Metabolite summary statistics

**Suppl. Table 3.** Summary statistics of the targeted short-chain fatty acid measurements

**Suppl. Table 4.** Associations between covariates and metabolites

**Suppl. Table 5.** Summary statistics of the comparison between eubiotic and dysbiotic IBD samples

**Suppl. Table 6.** Loads from the PCA based on metabolite levels

**Suppl. Table 7.** Summary statistics of the case–control analyses

**Suppl. Table 8.** Summary statistics of the case–control analyses of tryptophan and polyunsaturated fatty acid ratios

**Suppl. Table 9.** Summary statistics of the comparison between Crohn’s disease and UC metabolomics profiles

**Suppl. Table 10.** Relation between rare metabolites and medication use

**Suppl. Table 11.** Summary statistics of the association between metadata and the prevalence of metabolites

**Suppl. Table 12.** Summary statistics of the association between metadata and the levels of faecal metabolites

**Suppl. Table 13.** Associations between metabolite levels and disease characteristics

**Suppl. Table 14.** Summary statistics of the significant associations between host genetics and faecal metabolites

**Suppl. Table 15.** Conditional probabilities of co-occurrence between microbes and metabolites

**Suppl. Table 16.** Summary statistics of the association between metabolites and the presence of bacteria

**Suppl. Table 17.** Summary statistics of the association between metabolites and the abundance of bacteria

**Suppl. Table 18.** Summary statistics of the case–control analyses on microbial features

**Suppl. Table 19.** Summary statistics of the association between metabolites and the abundance of bacterial pathways

**Suppl. Table 20.** Summary statistics of the association between metabolites and the abundance of bacterial gene clusters

**Suppl. Table 21.** Summary statistics of the models predicting metabolites

**Suppl. Table 22.** Summary statistics of the models predicting IBD based on faecal metabolites

