## Supplemental Methods for "The faecal metabolome and its determinants in inflammatory bowel disease"

**SUPPLEMENTARY METHODS**

**Metabolite’s quantification**

Metabolomics measurements were performed by Metabolon Inc. (North Carolina, U.S.A.). In short, proteins and organic solvent were removed from each sample. Next, each sample was divided into four fractions for analysis: two for analysis by two separate reverse phases (RP)/UPLC-MS/MS methods with positive ion mode electrospray ionisation (ESI), one for analysis by RP/UPLC-MS/MS with negative ion mode ESI, one for analysis by HILIC/UPLC-MS/MS with negative ion mode ESI. Raw data processing and quality control were performed according to Metabolon's standards.

In addition to untargeted metabolomics, the concentration of eight short-chain and branched-chain fatty acids. i.e., acetic acid (C2), propionic acid (C3), isobutyric acid (C4), butyric acid (C4), 2-methyl-butyric acid (C5), isovaleric acid (C5), valeric acid (C5) and caproic acid (hexanoic acid, C6), were measured using LC-MS/MS methods. Acetic acid was the most abundant SCFA in the faecal samples (mean: 2339 μg/g, s.d. 1,131 μg/g), followed by butyrate (mean: 1,072.5 μg/g, s.d. 678 μg/g) and propionate (mean: 955.58 μg/g, s.d. 515.8 μg/g), while hexanoic acid presented the lowest concentrations (mean 74 μg/g, s.d. 110 μg/g).

**Phenotypes selection**

We retrieved the metadata including 180 entries consisting of dietary habits, medication and anthropomorphic measurements overlapping in both cohorts and 33 and 13 phenotypes specific to the IBD and the control cohorts, respectively. This information was included in further analyses if each category had at least 10 entries. A complete list of phenotypes is provided in **Supplementary Table 1**. Samples from patients with colectomy or stomas at the time of sample collection were removed from our analysis since their faecal samples are not representative of the content of the whole intestinal tract (n=68).

To adjust for differences in intestinal transit time, we combined the information about “bowel movements per day” present in the control cohort with questionnaires on the type of stools and frequencies a day in the IBD cohort. In the group of patients with IBD, disease remission or flare was defined using a combination of biomarkers, i.e., faecal calprotectin above 200 μg/g and Harvey-Bradshaw index>4 or Simple Clinical Colitis Activity Index (SCCAI) >2.5, and colonoscopy reports when available^1^.

Dietary intake was assessed through a validated food frequency questionnaire (FFQ) collected concurrently with faecal samples as described before^2,3^. Estimated food and nutrient intakes were adjusted by total caloric intake using regression analysis described in^4^. In addition, nutrient ratios and dietary patterns were calculated using pre-defined scoring systems:

- **Lifelines Protein score**, reflecting a higher protein energy percentage within the acceptable macronutrient distribution range for protein and a higher plant to animal protein ratio.
- **Lifelines Diet score,** expressing relative dietary quality with a higher score reflecting a high intake of vegetables, fruits, nuts, legumes and fish and lower intakes of red and processed meats and high sugar snacks and beverages.
- **Plant-to-Animal protein ratio**, reflecting a higher intake of plant protein relative to animal protein

**Metabolite ratios calculation**

In addition to individual metabolites, we calculated the ratios between molecules of interest. Ratios were calculated by dividing the raw metabolite’s levels, log transforming and scaling the resulting value.

In total, we evaluated 12 different ratios. The ratio between primary and secondary bile acids (deoxycholate/cholate, lithocholate/chenodeoxycholate, ursodeoxycholate/chenodeoxycholate), the ratio between conjugated and unconjugated bile acids (glycol + tauro bile acids / unconjugated bile acid), the ratios between kynurenine, tryptamine, serotonin and tryptophan and the ratios between omega-3 PUFA and omega-6 PUFA. In our dataset, we could quantify the levels of docosahexanoate (DHA), docopentaenoate (DPA), eicosapentaenoate (EPA), hexadecatrienoate and stearidonate as omega-3 PUFAs, and arachidonate, dihomolinoleate, dihomolinolenate, docosadienoate, hexadecadienoate and linoleate as omega-6 PUFAs.

**Prediction of microbial abundance**

Metagenomic reads mapping to the human genome were removed and reads containing Illumina adapters were trimmed using *KneadData* (v0.4.6.1)^5^. Other potential contaminants were also filtered out using *Kraken2*^6^ and the NCBI UniVec database, with the confidence parameter set to 0.5. After quality control of the sequenced reads, the microbial taxonomic and functional profiles were determined using *MetaPhlAn* (v3.0)^5^. Moreover, *HUMAnN 3.0* pipeline was used to estimate the metabolic potential of each microbial community^5^.

Three samples from patients with IBD were removed due to failure in the identification of bacterial species in their faecal sample. Previous to statistical tests, bacterial and pathway abundances were transformed using a centred-log ratio approach (CLR). Bacterial species and pathways present in more than 20% of the samples were kept for further analysis.

**Estimation of bacterial metabolic gene clusters in metagenomic samples**

Metagenomic reads were aligned to a collection of predicted metabolic gene clusters (MGC) predicted using GutSMASH^7^. BiG-MAP^8^ pipeline, with its default parameters, was used for read mapping and coverage calculations. In total, 6083 MGC were found in our dataset, for which, 1102 were kept after filtering for minimum coverage of 5% in the core genes of each cluster. To summarise the overall metabolic capacity of the microbial community, MGC were collapsed according to their predicted function by summing RPKM values. For example, the 5 different bai operons found in *Dorea sp. D27, Dorea sp AF36-15-AT, Clostridium scidens (ATC 35704), Clostridium hylemonae (DSM 15053)* and *Clostridium hiranonis (DSM 13275)* genomes, were merged into one *bai* operon category.

Centred log-ratio transformation was applied before data analysis. In total, 136 pathways were identified and 134 were kept for analysis after removing pathways that were present in less than 20% of the samples: “lysine degradation acetate to butyrate” and a “nitrate reductase”.

**Definition of dysbiosis**

Samples were defined as “dysbiotic” based on the microbiota composition in a similar way as described in Lloyd-Price et al.^9^. Euclidean distances between samples were computed on a clr-transformed bacterial abundances matrix. Non-IBD samples were used as a reference of eubiosis. Then, we computed the median distance between each sample and this reference group. A threshold for dysbiosis was defined at the 95^th^ quantile of the median distances between non-IBD samples. Samples exceeding this threshold were considered dysbiotic.

**Genome-wide association analysis power analysis**

Power estimations were conducted as described here^10^. First the relation between sample size and detection power was calculated while taking a grid search in the variance explained by the SNP (0.0~0.1). We then calculated the effects of metabolite detection rates (10%~100%) on the statistical power. The sample size in this study allowed us to have 80% power to detect genetic associations with 8% trait variation. The genetic effect of variants located in the NAT2 gene can explain 8% of the 5-acetylamino-6-amino-3-methyluracil variation (a metabolite with ~99% of prevalence in both IBD and controls) **(Suppl. Figure 6).**

**Defining host genetics combining whole-exome sequencing (WES) and global screening array (GSA)**

Library preparation, sequencing and variant calling were done at the Broad Institute of the Massachusetts Institute of Technology (MIT) and Harvard University. On average, 86.06 million high-quality reads were generated per sample and 98.85% of reads were aligned to a human reference genome (hg19). Moreover, 81% of the exonic regions were covered with a read depth >30x. Next, the Genome Analysis Toolkit (GATK) was used for variant calling^11^. Variants with a call rate <0.99 or Hardy-Weinberg equilibrium 𝜒2 test with p-value<0.0001 were excluded using PLINK 1.9^12^

GSA data was generated using the Infinium GSA-24 v1.0 BeadChip combined with the optional multi-Disease drop-in panel. Genotypes were called using OptiCall, QC steps were performed using PLINK 1.9 (variants with minor allele frequency (MAF) < 5%, call rate < 0.99 or Hardy-Weinberg equilibrium 𝜒^2^ test p-value<0.0001). Genotype data were phased using the Eagle^13^ and imputed to the Haplotype Reference Consortium reference panel using the Michigan Imputation Server^14^. After imputation, genetic variants were filtered for imputation quality R^2^ > 0.4. GSA genotype data was combined with WES data using PLINK 1.9. Variants with a MAF < 5% were removed.

In total, the combination of GSA and exome data covered 7,798,353 variants for 397 patients with IBD (CD =234 and UC=166) and 218 Lifelines Deep individuals.

**Prediction of IBD based on metabolomics profiles**

We used CoDaCoRe^16^ (v0.0.1) to identify ratios of metabolites and bacterial abundances that could predict IBD and its sub-phenotypes. Here, we first split the data into a training and a test set for each prediction, using 75% of the samples in the training process. Next, we estimated the added predictive value of using ratios of metabolites compared to a model including only host age, sex, BMI and faecal calprotectin levels (calprotectin levels >200 μg/g, yes/no). Patients with a history of intestinal surgeries (n=136) were excluded, and only highly prevalent metabolites (>70% of the samples) were considered in this analysis.

**Metabolite levels prediction**

For each of the metabolites and in each of the 8 defined models, we performed a 5-fold cross-validation (CV) procedure to select the best set of predictors based on the mean of squared errors. A 10-fold CV step was used in each of the CV-training sets to tune the lasso penalty parameter (lambda) in the lasso regression. Using the estimates of the model minimising the mean of squared errors, we computed the R^2^ coefficient in the whole dataset. We defined 8 different models representing different data categories available in both cohorts (IBD and non-IBD samples).

1) Host and technical factors: Which included information about the sex, age, BMI, average bowel movements per day, storage time at -80°C, batch and amount in grams of sample used for measuring metabolomics. All other models also included these variables to consider confounders' effects.
2) Diet: 119 dietary food patterns adjusted by total caloric intake.
3) Biomarkers: The levels of chromogranin A, human-beta defensin 2 and faecal calprotectin levels above 200 (yes/no).
4) Medication: The use of 22 medication categories (yes/no).
5) Disease: IBD (yes/no)
6) Taxa abundance: Relative abundance of 109 microbial species.
7) Bacterial pathways: 326 MetaCyc pathways
8) All: A model containing all variables described in the previous model.
